# Three-step scalable production of extracellular vesicles from pancreatic beta cells in stirred tank bioreactors promotes cell maturation and release of ectosomes with preserved immunomodulatory properties

**DOI:** 10.1101/2024.09.05.611247

**Authors:** Laurence de Beaurepaire, Thibaud Dauphin, Quentin Pruvost, Apolline Salama, Aurélien Dupont, Laurence Dubreil, Dominique Jégou, Grégoire Mignot, Benjamin Mahieu, Julie Hervé, Blandine Lieubeau, Jean-Marie Bach, Steffi Bosch, Mathilde Mosser

## Abstract

Small extracellular vesicles (sEV) released by healthy beta cells are promising candidates for diabetes therapy thanks to their aptitude to modulate inflammation, to induce or maintain pancreatic function and to prevent pathogenic mechanisms. To advance the clinical development of therapeutics, there is a crucial need for scalable production methods. Stirred tank bioreactors (STR) are widely used in the industry due to their ability to provide homogeneous gas and nutrient supply, online monitoring, and efficient scale up. Anchorage-dependent cells can be cultured in STR on microcarriers or as spheroids, but may experience shear stress, which can affect sEV phenotype and function. Using pancreatic beta cells, this study identifies critical cell culturing parameters, including culture mode (monolayer vs. spheroids), medium formulation (with or without serum, glucose control), and process parameters (stirring, duration, cell density). The findings show that small spheroid culture promotes beta cell maturation without decreasing the yield of sEV per cell, despite a reduced cell surface exchange area. However, stirring increased expression of cellular stress markers and decreased cell viability. Set up of a three-step bioprocess allowed to maximize cell viability and sEV yields at high cell density over short production duration. sEV produced under these conditions maintained high purity, membrane integrity, and the aptitude to reduce T-lymphocyte proliferation and IFN-γ cytokine secretion in a mixed lymphocyte reaction. Flow cytometry analysis revealed lower CD63/CD81 ratios in STR, indicating enhanced ectosome production. Switch from high glucose expansion to low glucose production medium further allowed to direct sorting of the antigen insulin into beta-sEV. This study demonstrates the feasibility of producing functional sEV from mature beta cells cultured as small spheroids, suitable for upscale. Production of sEV in STR may be particularly beneficial for ectosome-enriched compound loading for therapeutic applications.

**Graphical abstract:** Rational development of a scalable bioprocess to control extracellular vesicles production & function.

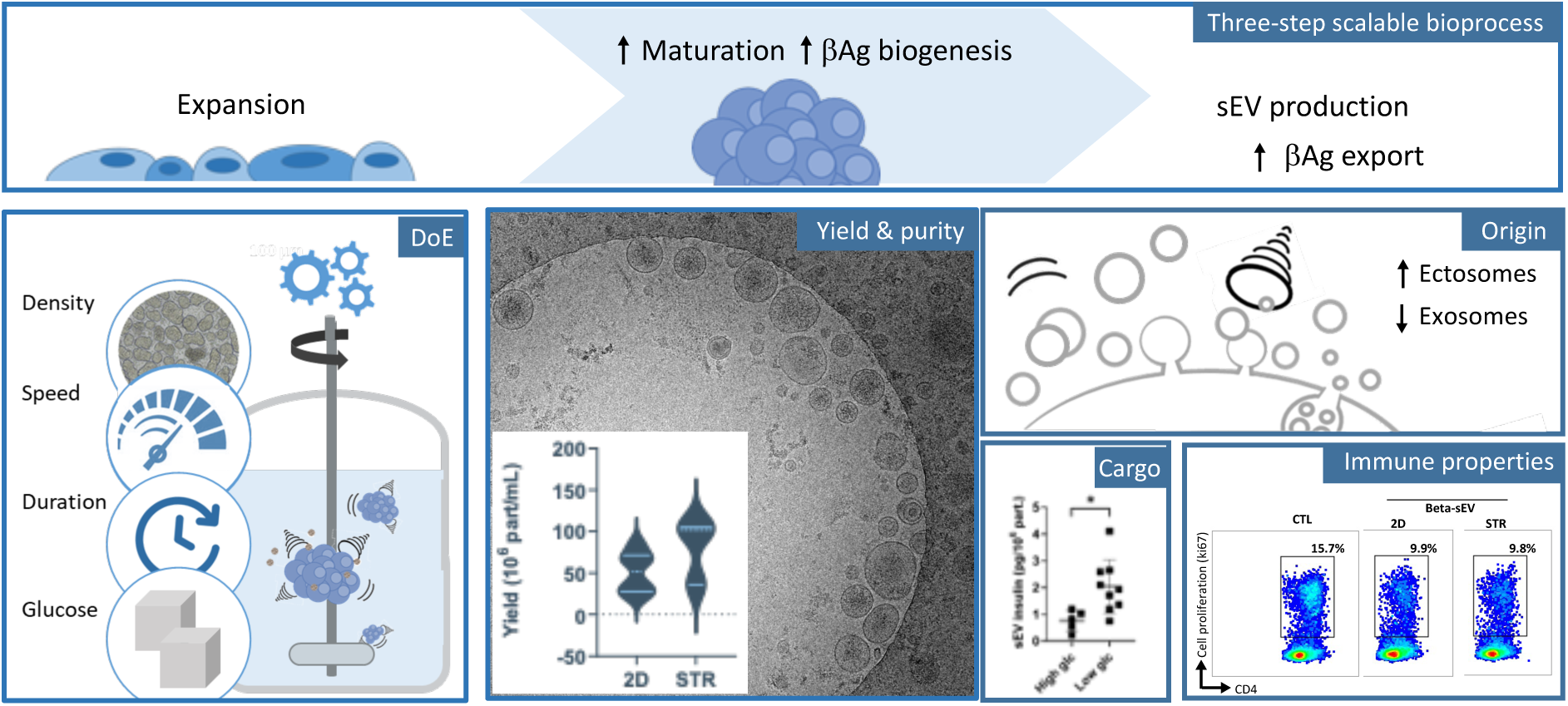

**Highlights:**

- Small 3D-spheroid culture promotes cell maturation
- Stirring induces cellular stress responses & promotes ectosome release from the cytoplasmic membrane
- Design of experiment efficiently enhances cell viability and EV yield with preserved immune-modulatory properties
- Glucose-starvation during the production phase directs insulin-sorting into extracellular vesicles

## Introduction

Type 1 diabetes mellitus (T1D) is a chronic disease that results from the autoimmune destruction of insulin-secreting beta cells within the islets of Langerhans in the endocrine pancreas. The disease affects more than nine million people worldwide (WHO, https://www.who.int/news-room/fact-sheets/detail/diabetes) with an annual incidence rate growing by 3% per year [1] and a cost of medication reaching $100 billion worldwide (www.idf.org). Despite notable improvements in diabetes care, no cure exists to date and patients experience serious long-term complications and reduced life expectancy (www.t1dindex.org). Ideally, reprogramming of autoimmunity towards tolerance would allow to prevent disease progression and to preserve the residual pancreatic beta cell function at diagnosis and the long-term engraftment of pancreatic islets in transplant recipients [2,3]. Technically, the induction of protective immune memory requires the presentation of the specific antigen(s) in the presence of appropriate tolerogenic signals. Major challenges concern the selection of target antigens, their formulation, dose and route of administration [3,4]. In T1D patients, diversification of the autoreactive immune lymphocyte repertoire during the asymptotic phase preceding diagnosis jeopardizes mono-antigenic approaches used in earlier clinical trials (DiaPep277, Diamyd, oral insulin, reviewed in [5]).

Small Extracellular vesicles (sEV) released by living cells are promising candidates for multi-target therapies. Highly abundant in body fluids, fully biocompatible, EV act as natural vectors of cell-to-cell communication that have supplanted synthetic vehicle formulations in their ability to cross even difficult biobarriers and deliver cargo to distant recipient cells [6,7]. They display a propensity to create an immunosuppressive environment in health and disease [8–10]. In comparison to cell therapy products, sEV are non-replicative, filter-sterilisable, storable off-the-shelf products with reduced sanitary risks [11,12]. *In vivo*, neither the transfer of high titers of allogeneic EV during blood transfusions, nor the repeated administration of allo- or xenogenic EV in immuno-competent animals and clinical trials engender notable adverse events [7,13,14]. In addition to the well-documented role of stem cell EV, evidence emerges on contributions of EV from mature healthy tissue to immune homeostasis [10,15]. sEV derived from pancreatic beta cells (beta-sEV), convey a cocktail of beta antigens challenging to produce and stabilise in their native conformation e.g. (pro-) insulin (INS), glutamic acid decarboxylase (GAD65), zinc transporter 8 (ZnT8) and islet antigen-2 (IA2) [16–19]. Beta-sEV participate in protective and adaptive mechanisms in the beta cell in physiological situations [20]. To date, murine and human allogeneic beta-sEV have been shown to improve the function, viability and immune protection of pancreatic beta cells [21–23].

sEV are complex polydisperse bioproducts that mirror the state of their parental cell. Evidence accumulates that culture conditions are decisive for versatile properties of the sEV produced [24]. We and others reported earlier that exposure to stress in culture mirroring pathological conditions such as inflammation and hypoxia [17,20,25] modifies the release and phenotype of beta-sEV. Pancreatic beta cells are highly performant secretory cells capable to adapt quickly to changes in food intake or energy demands raised by physical activity or inflammation [26,27]. However, its demanding secretory function renders the beta cell vulnerable to stress. Indeed, massive induction of insulin synthesis can exceed the beta cell’s endoplasmic reticulum (ER) folding capacity, leading to the accumulation of misfolded proteins and subsequent activation of stress signalling pathways to restore homeostasis [27]. The culture conditions are therefore decisive for the properties of the sEV produced, in particular from stress-sensitive parental cells.

For (pre-) clinical testing, the use of a continuous cell line is essential to produce large amounts of allogeneic beta-sEV. Indeed, the production of beta-sEV from primary cells or differentiated stem cells is severely limited in quantity, standardization and safety. Pancreatic beta cells are anchorage-dependent cells that spontaneously self-aggregate to form spheroids - hereafter named pseudo-islets because of their resemblance to pancreatic islets - in suspension culture vessels [28]. This three-dimensional organization has been shown to promote beta cell maturation [29]. Based on the average yield of 2.5 µg EV protein/10^6^ producer cells, calculated from the results of 54 publications [30] and in accordance with our studies on beta cell EV [25,31], the treatment of a 70 kg human would require the production of EV by ∼2.6×10^11^ cells. In line with these estimations, 10^8^ to 10^12^ particles have been administered per patient in recent clinical studies [32,33]. Almost exclusively produced at the laboratory scale, these applications demand a critical upscaling of the EV manufacturing process.

The majority of studies produced sEV from adherent cells cultured in standard flasks inappropriate for large-scale production. Static multi-chamber bioreactors e.g. the CeLLine® bioreactor could potentially improve sEV production yields by a factor of 10 to 100 [34–36], but this system is still limited to a volume of 1 L. Other studies have successfully used hollow fibre bioreactors with a culture area of up to 2.1 m^2^ [37,38]. Cells grown at high density form a 3D network between fibres, while sEV accumulating in the extra-capillary space within the hollow fibres can be repeatedly harvested. Alternatively, stirred tank reactors (STR) allow to culture adherent cells on biomaterials (hollow fibres, microcarriers, microcapsules) or as spheroids in volumes of up to 2,000 L (eq. 1,080 m^2^)[39,40]. STR are most commonly used in the industry because of their proven performance [41]. Among notable advantages, they present reduced risks of contamination, ease of scale up, and maintenance of homogenous conditions by mixing for continuous gas and nutrient supply. At/off line monitoring of pH, temperature, glucose consumption, cell growth & death allow to evaluate precisely the parental cell’s status during the upstream phase and to adjust culture feed perfusion over time.

STR have been successfully used for the production of sEV from cell lines including HEK293 and MSC [42–45]. While several studies reveal increased sEV yields in comparison to static 2D cultures, they also report phenotypic changes of the final product with functional implications. In addition to the cell source, major parameters to consider for sEV production in STR are cell density at seeding, duration of culture, speed of stirring, medium formulation and interactions in between. Hydrodynamic forces engendered by excessive agitation can improve EV yields, but can be source of harmful cell stress and death [33,46,47]. Furthermore, sEV production media should be tested EV-free, ideally, chemically defined, all while supporting the growth and/or maturation of the parental cells.

The aim of the present study is to develop a scalable process to produce large quantities of highly pure beta-sEV with a preserved “healthy” phenotype for diabetes therapy. For process development, one-factor-at-a-time analyses do not inform about parameter interactions. Design of experiments (DoE) tools combine variance and pairwise interaction analysis allowing fine-tuning of parameters with high efficacy and timeliness [48]. Using a continuous beta cell line grown as pseudo-islets in a STR, we first screened for the impact of upstream process change on the beta cell performance for sEV production. Parameters studied included the medium formulation (w/ or w/o serum, glucose refeed), the mode of culture (monolayer, spheroids, static, stirred) and the process parameters (stirring speed, duration, cell density). We assessed the impact of these parameters on beta cell maturation, viability, reticulum stress and morphology as well as on sEV production yields. These results allowed the development of a three-step scalable process with improved process parameters. Second, the performance of this process was characterized with regard to relevant sEV quality attributes such as their morphology, size, purity, expression of EV markers, insulin content and immune activity. Our results demonstrate that beta-sEV are most efficiently produced in STR in serum-free medium seeded at high cell density over short production times. Stirring, even at speed below shear stress thresholds, further promotes release of ectosomes with a slightly increased size, but preserved immune-modulatory function. Lowering glucose concentration during the production phase also enhances the direct sorting of the beta antigen insulin into sEV. STR bioprocess solutions for mature, stress-sensitive cells unlock a new spectrum of sEV therapeutic application in the future, especially for the loading of cytosolic and cytoplasmic membrane components for immune therapy. In perspective, the development of optimised animal-component-free EV production media is required to increase further sEV productivity in perfusion culture.

## Materials and methods

### Beta cell culture

MIN6 cells (kindly provided by Prof. Jun-ichi Miyazaki, University Medical School, Osaka, Japan) were cultured as previously described [25]. Briefly, for routine expansion, MIN6 cells were grown as adherent cells in tissue culture flasks –T-flask) plated at a density of 1.5 x 10^5^ cells/cm^2^ in DMEM high glucose (4.5 g/L) medium (Life Technologies, Saint Aubin, France) supplemented with 10% Fetal Calf Serum (FCS, Eurobio, Les Ulis, France) and 20 µM beta-mercaptoethanol (SIGMA, Saint Quentin Fallavier, France). Cultures were regularly assessed for mycoplasma contamination using the MycoAlert TM mycoplasma detection kit (Lonza, Basel, Switzerland). For pseudo-islet formation under static or dynamic conditions, MIN6 cells were seeded in complete medium at a density of 1×10^6^ cells/mL onto non-surface coated petri dishes or spinner (Corning, Sigma Aldrich) at a stirring speed set to 90 rpm (Biomix, 2mag, Munich, Germany). Prior to EV production, MIN6 cells were washed twice in PBS w/o Ca^2+^ and Mg^2+^ (Eurobio) and switched to serum-free OptiMEM low glucose (1 g/L) medium (Fisher Scientific, Illkirch, France) supplemented with 20 µM beta-mercaptoethanol at densities and glucose concentrations as indicated in figure captions. Conditioned supernatants were harvested 4 h, 24 h or 44 h later for EV isolation.

### Experimental design

The impact of cell culture conditions on MIN6 beta cells were evaluated using a fractional factorial design, focusing on the main factors and their first-order interactions. In a first series of experiments, the three factors studied were culture mode (monolayer/pseudo-islets), agitation (static/ stirred) and medium formulation (w/ or w/o FBS). The stirred monolayers (w/ or w/o FBS) were excluded as they are not considered relevant culture conditions. The response variables measured were *(i)* cell viability, determined by counting viable cells after trypsin dissociation and trypan blue staining, *(ii)* dead and lysed cells, assessed by measuring lactate dehydrogenase (LDH) activity in cell supernatant and *(iii)* relative *C/EBP homologous protein (*CHOP) expression compared to the reference culture (monolayer in medium w/ FBS). Initially, MIN6 beta cells were cultured as monolayer in T-flasks or as static pseudo-islets in Petri dishes (PISTAT) or in stirred spinners (PISTIR) for three days in medium w/FBS. Afterward, the medium was replaced, and MIN6 cells were cultured as monolayer, static or stirred pseudo-islets, either in medium w/FBS or w/o FBS for 24 h at a cell concentration of 2.5 ×10^6^ cells/mL. Subsequently, a full factorial design with central point was warried out aiming to fine-tune the process parameters during sEV production in order to maximize the cell viability, to minimize cellular stress and to maximize sEV production by tuning the stirring, the cell density and the duration of culture. The three factors studied were stirring (60, 90 and 120 rpm), cell density (1, 2.5 and 5 x 10^6^ cell/mL) and duration (4, 24 and 44 hours) of culture. The response variables were cell viability (%), LDH activity, CHOP relative expression compared to PI cultured in a stirred system at 90 rpm in medium w/ FBS and sEV concentration (particle/mL) and sEV production (particle/cell).

### Assessment of cell viability

Cell viability was evaluated using trypan blue exclusion dye (Fisher Scientific) on trypsin dissociated cells. As a marker of cell death, lactate dehydrogenase (LDH) activity was measured in culture supernatants using a colorimetric detection kit (Roche, Diagostics, Meylan, France).

### Glucose-stimulated insulin-secretion assay

To evaluate beta cell function of MIN6 monolayers or pseudo-islets, 10^6^ cells were pre-incubated in a P6-well in low glucose medium composed of RPMI supplemented with 2 mM L-glutamine, 2.8 mM glucose and 0.5% bovine albumin for 60 min at 37 °C, 5% CO2. Following this period, cells were stimulated at high 20 mM glucose concentration in the presence of 10 mM theophylline for one hour. Culture supernatants were then carefully sampled and centrifuged at 200 g for 3 min to avoid accidental islet uptake and stored at -20°C until analysis. Total mouse insulin secreted by MIN6 cells was quantified by ELISA (Mercodia, Upsala, Sweden). Stimulation indexes of insulin secretion in response to glucose stimulation is calculated by the ratio of insulin after high glucose plus theophylline stimulation over basal insulin secretion release (mM/mM).

### Total protein content

For protein analyses, cells were lysed in RIPA lysis buffer containing proteases/phosphatases inhibitors (Ozyme, Saint Cyr l’Ecole, France) or in ethanol acid buffer followed by Tris 1M pH 7.5 neutralization. sEV were lysed in 0.1% Triton X-100 (SIGMA Aldrich). Protein concentrations were determined by a Bradford protein assay using Coomassie Plus assay reagent (Fisher Scientific) following the supplier’s recommendations. Optical densities were read on a Fluostar instrument (BMG LABTECH, Champigny sur Marne, France).

### Insulin content

For quantification of intracellular insulin content, 1-5×10^6^ MIN6 cells grown as monolayers or pseudo-islets were lysed in 125 µL acidic ethanol buffer (1.5% HCL in 70% EtOH) for insulin extraction and stored at – 80°C until quantification by ELISA. For sEV insulin quantification, vesicles were lysed by addition of 0.1% triton X-100 and 2 min of ultra-sound exposure prior to ELISA analysis according to the supplier’s recommendation (Mercodia).

### Transcriptomic analysis

Total RNA was extracted from MIN6 cells using the miRVana extraction reagent (Fisher Scientific). Prior to quantitative real-time PCR analysis, 3 µg of RNA were treated with TurboDNase (Fisher Scientific) and reverse transcribed using M-MLV Reverse Transcriptase (Fisher Scientific) and random 15-mers primers (Eurogentec, Angers, France). Mouse markers of beta function, endoplasmic reticulum stress and EV biogenesis pathways were quantified using primer sequences described earlier [49–51] and Evagreen reagent (Solisbiodyne, Tartu, Estonia) on a CFX96 Touch Real-time PCR detection system (Bio-Rad, Marnes La Coquette, France) in application of the relative standard curve method [52]. Relative quantities of target gene expression were normalised with the geometric mean of the sample’s housekeeping genes GAPDH and HPRT1 expression.

### Isolation of beta sEV

EV were collected from MIN6 supernatants using a method combining differential centrifugation, tangential flow filtration and size-exclusion chromatography. Briefly, conditioned culture media was centrifuged immediately after harvest at 300 x *g* 10 min, 2,000 x g 20 min and 16,000 x g, 20 min to discard cell debris and large vesicles. The 16,000 x g supernatants were filtered 0.2 µm and sEV isolated using a 300 kD MidiKros column (Repligen, Breda, Netherlands) on a KrosFlo Research IIi Tangential Flow Filtration (TFF) instrument (Repligen). The input flow rate was set to 50 mL/min resulting in a transmembrane pressure of approximately 300 mbars. TFF comprised four cycles of diafiltration before final concentration down to 20 mL. Whenever indicated, sEV were concentrated down to approximately 100 µL on an AMICON MWCO-100 kDa cellulose ultrafiltration unit (Fisher Scientific) and passed through a qEV 35 nm single size exclusion chromatography column (IZON, Lyon, France). sEV were collected in flow-through fractions two and three. sEV were stored at 4 °C for one to three days or at -80 °C in PBS 25 mM Trehalose [31]. Whenever indicated, sEV were precipitated using ExoQuick exosome isolation reagent (Ozyme, Saint Cyr l’Ecole, France) from 2.5 or 5.0 mL of crude culture supernatant and processed following the supplier’s recommendation.

### Tunable resistive pulse sensing (TRPS)

The size and concentration of sEV were analysed by TRPS using a qNANO instrument and NP100 or NP150 nanopores (IZON). All samples were diluted in PBS 0.03% Tween-20 electrolyte. After instrument calibration with 110 nm or 210 nm calibration beads (Izon) for UF/TFF or Exoquick purification-based methods, respectively, all samples were analysed with recording at least two different pressures.

### sEV flow cytometry bead assay

Magnetic streptavidin beads (4.5 µm; Fisher Scientific) were coated with 4 µg of biotinylated anti-mouse CD81 capture antibody (clone Eat-2) per 10^7^ beads for one hour at room temperature in PBS 0.1% BSA. After washing, sEV (3 x 10^4^ particles/bead) were pulled down by over-night incubation with beads at 800 rpm 4 °C. The following day, beads were washed and stained with PE-CD63 (clone NVG-2; 2 µg/mL), APC- CD81 (clone Eat-2; 2 µg/mL) detection antibodies or isotypic controls (all antibodies were purchased from BioLegend). Flow cytometry was performed on a MACSQuant 10 instrument (Miltenyi Biotech, France). Single beads were gated for analysis of mean fluorescence intensity (MFI). Tetraspanin surface ratios were calculated as follows: (MFI CD63 – MFI CD63 isotypic control) / (MFI CD81 – MFI CD81 isotypic control).

### sEV immunofluorescent tetraspanin staining

Exoview mouse tetraspanin kit (EV-TETRA-M2, NanoView Biosciences) was used for sEV tetraspanin detection. According to the manufacturer’s protocol, sEV were captured by anti-CD81 antibody on a chip and detected by immunofluorescence with anti-CD81 (green), anti-CD63 (red) and anti-CD9 (blue) labelled antibodies provided by the kit on an Exoview R200 instrument. Data were analysed using Exoviewer 3 software for particle count and co-localisation analysis of individual capture spots.

### Cryo-electron microscopy

For morphological analysis, sEV were prepared for cryo-electron microscopy as described earlier [25] and imaged on a Tecnai G^2^T20 sphera electron microscope (FEI company, The Netherlands) equipped with a CMOS camera (XF416, TVIPS) at 200 kV. Micrographs were acquired under low electron doses using the camera in binning mode 1 and at a nominal magnification of x 25,000. sEV size measurements were performed manually using Fiji software (version 1.54).

### Confocal Imaging

For immunofluorescence staining, 2x 10^5^ MIN6 cells per well were seeded onto 8-chamber Nunc LabTek slides. The following day, cells were washed in PBS and fixed with PBS 4% paraformaldehyde for 15 min at room temperature before permeabilisation with PBS 0.3% Triton X-100, 5% bovine serum albumin for 15 min at room temperature. Cells were washed again and incubated with PBS 5% BSA blocking buffer for 30 min at room temperature, followed by overnight incubation at 4 °C with primary antibodies or isotypic controls diluted 1:100 in blocking buffer (anti-mouse CD63, clone NVG-2; APC-anti-mouse/rat CD81, clone Eat- 2, and anti-mouse CD9, clone MZ3). The next day, cells were washed again and, for CD63 and CD9 detection, incubated with a secondary AF488- goat anti-rat IgG2k antibody for one hour at room temperature. For insulin staining, monolayers or pseudo-islets included into Corning^TM^ Matrigel^TM^ Matric (Fisher Scientific) and sliced, were saturated with PBS 5 % donkey serum, prior to incubation with primary rabbit anti-insulin antibody (clone C27C9, 1:400; Ozyme) overnight at 4 °C. The following day, slides were washed with PBS and incubated with AF488-donkey anti-rabbit antibody (1:2000) (Fisher Scientific) for one hour. Nuclei were counterstained with Hoechst 33342 (SIGMA) at 1 µg/mL for 10 min. at 37 °C before imaging on a LSM780 confocal microscope (Zeiss, Oberkochen, Germany) and analysis with Fiji software [53].

### Mixed lymphocyte reaction

To assess for immuno-regulatory abilities of beta sEV, a mixture of cells spleens isolated from 3-4 inbred mouse strains (BALB/c, C57BL/6, FVB, C3H) purchased from Charles River (Saint Germain Nuelles, France) was seeded per well onto round-bottomed P96 well plate as follow : 4 x 10^5^ total cells per well of equal proportion of each mouse strain in 200 µL of X-VIVO-15 medium (Lonza), supplemented with 10 µg/mL transferrin (Lonza), 100 IU/mL penicillin, 100 µg/mL streptomycin, and treated with 7 x 10^8^, 2 x 10^9^, or 6 x 10^9^ of sEV (particles/well/200 µL). Controls (final working concentration) included were vehicle only PBS trehalose (SIGMA; 2.5 mM) and rapamycin (Miltenyi, Paris, France; 100 nM). Plates were centrifuged 300 x g, 1 min to favour encounter of immune cells prior to culture at 37 °C in a 5% CO2 humid atmosphere. After four days of culture, cells were harvested and analysed by flow cytometry for expression of the Ki67 proliferation marker (clone REA, dilution 1:50; Miltenyi) using a fixable viability dye (FVD-450, Fisher Scientific) and intracellular staining reagents (Fisher Scientific) according to the supplier’s recommendations. Culture supernatants were collected and processed for cytokine analysis using a mouse Th cytokine panel (12-plex) cytometric bead assay (BioLegend, Amsterdam, Netherlands). Flow cytometry was performed on a MACSQuant II Flow cytometry instrument (Miltenyi). Data were analysed using FlowJo v10.8.1 software (Tree Star Inc., Ashland, OR, USA).

### Statistical analyses

Analysis of variance (ANOVA) and regression analysis were performed using the Statgraphics Centurion software 18.1.06 (Francestat, Neuilly sur Seine, France). The significance of differences between groups was evaluated using statistical tests as indicated in figure captions. A p-value <0.05 was considered statistically significant. Graph formatting was performed using Graphpad Prism software 9.5.1 (Comparex, Issy-les-Moulineaux, France).

## Results

### Upstream strategy to produce sEV from healthy beta cells at a large-scale

The usefulness of beta sEV to modulate diabetogenic responses is further licensed by the beta cargo and function inherited from their cell of origin. Consequently, upstream culture conditions have to warrant healthy beta cell growth and maturation. Cultivated on hydrophobic surfaces, MIN6 beta cell spontaneously formed small 3D spheroids suitable for culture in STR in the absence of scaffolds or polymers (**Figure 1A**) with a mean size ± SEM of 59 ± 0.76 µm, staining positive for insulin (**Figure 1B-C**). This organization extended their population doubling time by 34 % (p<0.05), from 33.6 ± 2.6 h to 45.0 ± 3.1 h (**Figure 1D)**, while enhancing their ability to produce insulin in response to glucose stimulation by 33 % (p<0.05), with a stimulation index increasing from 2.28 ± 0.21 to 3.04 ± 0.26 (**Figure 1E)** for cells grown as monolayers or stirred pseudo-islets, respectively. Enhanced beta cell maturation is further supported by an 1.9 fold increase of the homeobox protein Nkx6.1 (Nkx6.1) transcription factor (p<0.01) and an 1.8 fold increase of the inducible insulin 1 (INS1) gene expression (p<0.05) (**Figure 1F**). 3D culture did not significantly alter expression of the pancreatic duodenal homeobox 1 (PDX1) and insulin 2 (INS2) genes. According to the standard culture protocol for MIN6 cells [54] and our observations of beneficial effects of glucose on MIN6 proliferation and insulin biosynthesis (**Suppl. Figure 1**), these experiments were performed in high glucose (4.5 g/L) DMEM medium. These observations led to the set-up of a three-step upstream strategy comprising a phase of beta cell expansion as 2D monolayers followed by a phase of maturation via the formation of pseudo-islets in STR for sEV production.

**Figure 1.**
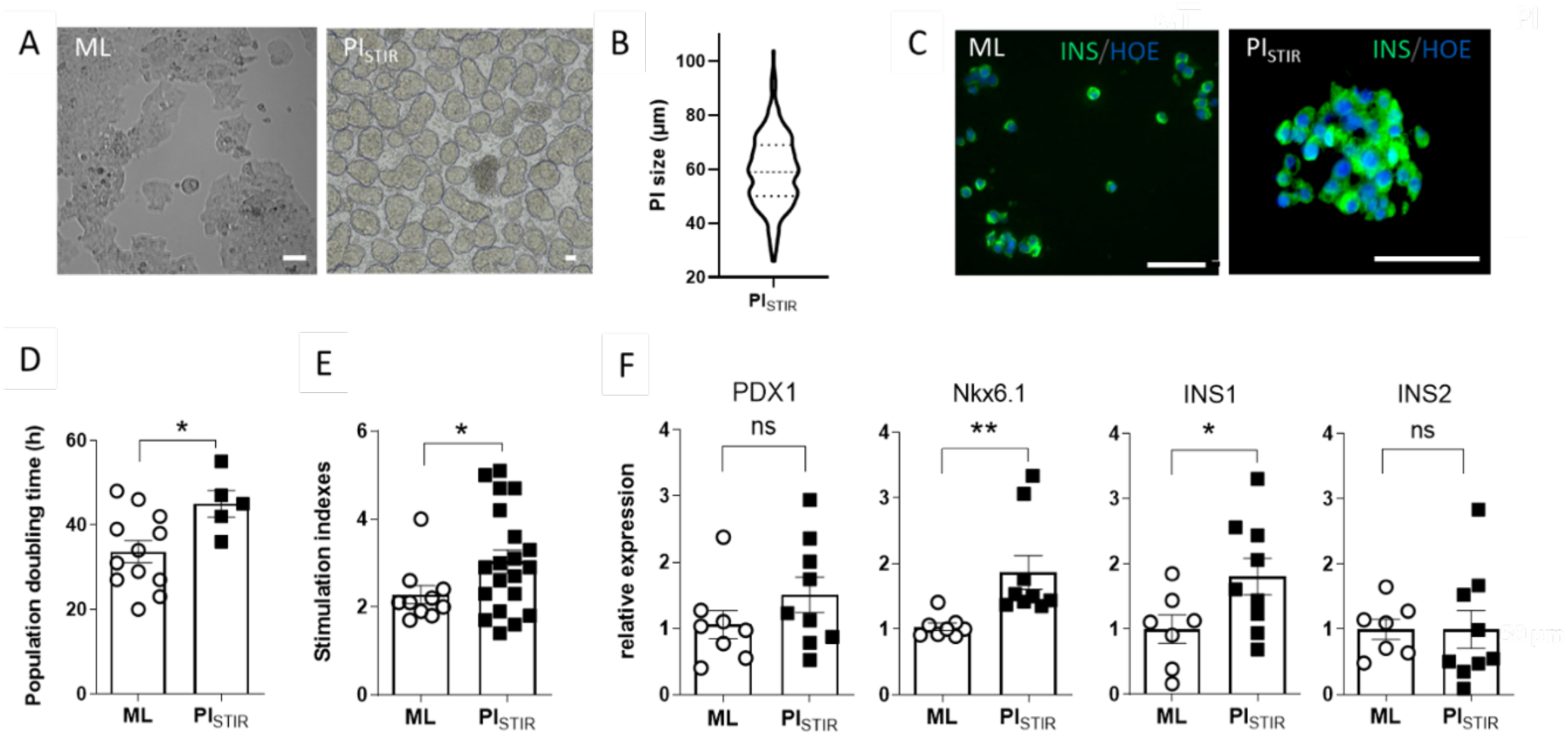
3D spheroid culture promotes beta cell function. MIN6 beta cells were cultured as monolayer (ML) in T-flasks or as pseudo-islets (PI) in spinners at 90 rpm for three days. (A) Representative microscopy images. Scale bar: 100 µm. (B) Size distribution of PI measured manually using Fiji software. Representative violin plot with median and quartiles of one out of three independent experiments (n=304). (C) Immunofluorescence staining of insulin (in green) in MIN6 cells with nuclei counterstained with Hoechst (in blue). Representative microscopy images. Scale bar: 50 µm. (D) Population doubling time, (E) Stimulation indexes of insulin secretion in response to glucose stimulation, calculated by the ratio of insulin secretion after high glucose-theophylline stimulation over basal insulin secretion release (mM/mM), and (F) Transcriptomic analysis of beta cell function by real-time RT-PCR represented as fold change relative to ML. (D-F) Results are depicted as individual values and means ± SEM from at least five independent experiments. Unpaired parametric t- test (*p<0.05, **p<0.01).

Clinical grade sEV production requires use of animal component-free, preferentially chemically defined, culture reagents. With the aim to produce sEV from healthy beta cells, we next characterized the impact of static vs stirred culture conditions in serum-free medium on beta cell viability and cellular stress pathways. The impact of the following three factors was evaluated on the health status of the cells using a fractional factorial design: mode of culture (monolayer vs pseudo-islet), agitation (static vs stirred systems) and medium formulation (w/ or w/o FBS) (**Figure 2** and **Supp Table 1**). The stirring of pseudo-islets (**Table 1**, **Figure 2A-C**) tended to decrease cell viability (p=0.0684) and significantly increased CHOP expression (**Figure 2D**, p<0.0001), while no significant impact was observed for solely serum deprivation or 3D culture mode. Both culture mode and serum deprivation (alone or in interaction) significantly increased LDH release in culture supernatant (**Figure 2C**, p<0.05). In contrast, stirring alone did not engender LDH release. Finally, the concentration of sEV produced in the conditions of culture w/o FBS was quantified after precipitation. A significant (> 4-fold) increase in the amount of sEV produced from pseudo-islets was observed in stirred systems (p<0.001), while no difference was observed between sEV produced by pseudo-islets cultured in static conditions (**Figure 2E**). These results underlined the feasibility to produce sEV from beta cells cultured as pseudo-islets in serum-free medium and that agitation promotes sEV yield, but with a negative impact on cell viability and stress demanding further process optimisation.

**Figure 2.**
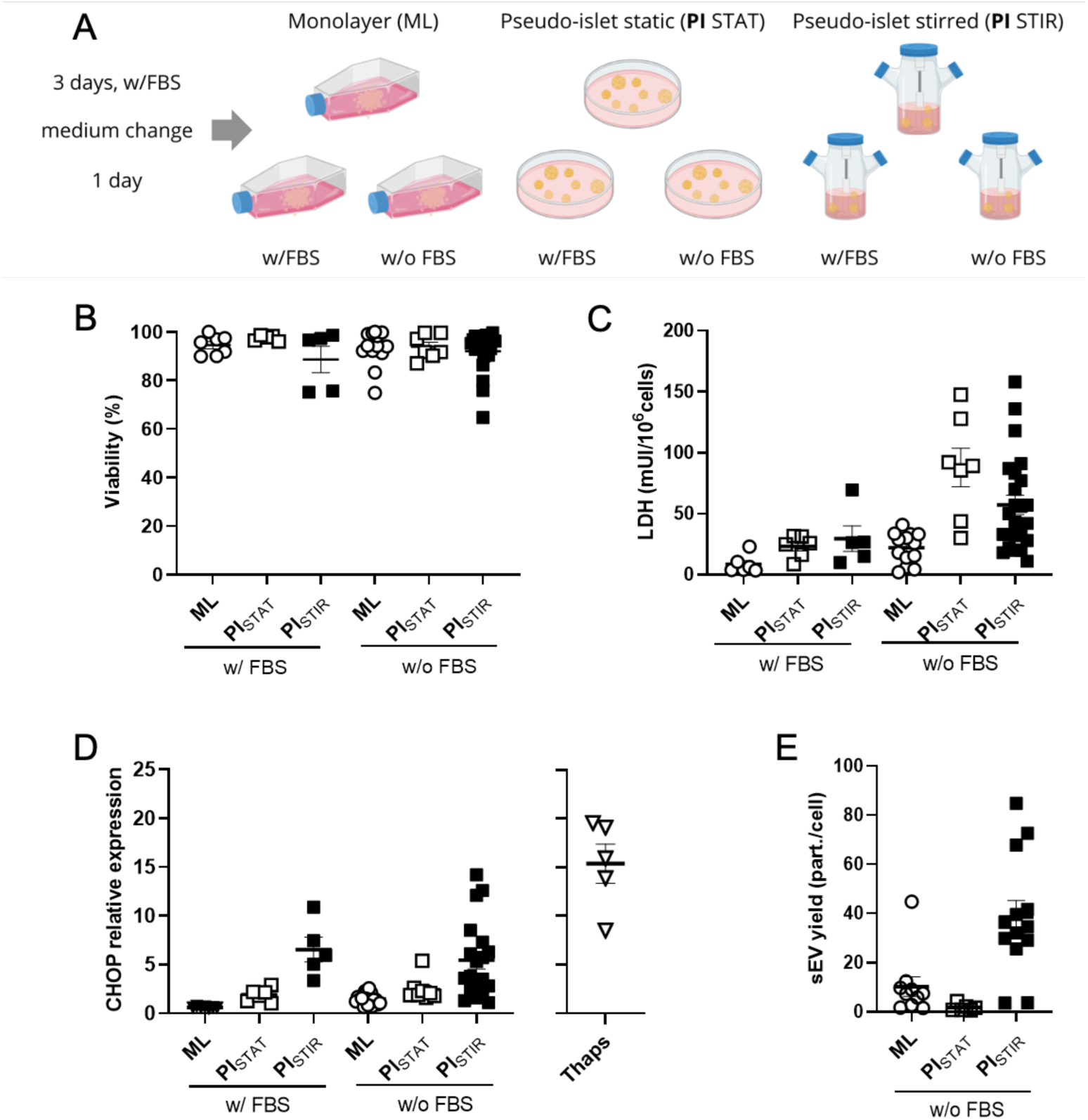
Stirring increases cell stress and sEV production. (A) MIN6 beta cells were cultured as ML in T-flasks or as static PI in Petri dishes (PI_STAT_) or in stirred spinners (PI_STIR_) for three days in medium w/FBS. Then, the medium was changed and MIN6 cells were cultured as ML, static or stirred PI either in medium w/FBS or w/o FBS for 24 h at a cell concentration of 2.5 ×10^6^ cells/mL. (B) Cell viability was assessed by measuring the total of viable cells after trypsin dissociation and trypan blue staining, and (C) Cell death by measuring the LDH activity in culture supernatants (D) CHOP expression represented as fold change relative to MIN6 cells grown as ML w/FBS. MIN6 cells treated with 1 µM of thapsigargin (Thaps), an ER stress inducer, for 24 h in medium w/FBS were included as a positive control. (E) The yield of sEV per cell isolated from culture supernatants by the precipitation method determined by TRPS. Results are depicted as individual values of biological replicates and means ± SEM from at least five independent experiments.

**Table 1.** Estimated effects of factors, their interactions and the statistical significance thereof to identify culture parameter impact on beta cell performance for sEV production.

| Factors | Viability (%) |  | LDH (mUI/10<sup>6</sup> cells) |  | CHOP (relative expression) |  | sEV concentration (10<sup>6</sup> part./mL) |  |
| --- | --- | --- | --- | --- | --- | --- | --- | --- |
|  | effect | p-value | effect | p-value | effect | p-value | effect | p-value |
| Hydrodynamic : Stat versus Stirred | -2.48 | 0.0684 | -6.08 | 0.2822 | 1.9651 | **0.0001** | 21.7 | **<0.0001** |
| Culture mode : ML versus PI | -0.84 | 0.5245 | 20.15 | **0.0007** | 0.5633 | 0.1907 | -4.27 | 0.2900 |
| Medium : w/ FBS versus w/o FBS | -0.71 | 0.5530 | 10.29 | 0.0557 | 0.0096 | 0.9811 | / | / |
| Culture mode x medium | -0.58 | 0.6612 | 12.85 | **0.0258** | 0.0795 | 0.8524 | / | / |
| Medium x hydrodynamic | 1.92 | 0.1549 | 9.29 | 0.1032 | 0.3449 | 0.4413 | / | / |
| **R<sup>2</sup>** | 6.6 % |  | 41.2% |  | 46.5 % |  | 63.0 % |  |

\*p-values computed by StatGraphic software from the analysis of variance performed for each response reaching statistical significance $\geq 5\%$ are highlighted in **bold**

As stirring is essential to scale up the process in large-scale STRs, we carried out a full factorial design with a central point aiming to fine-tune the process parameters in order to mitigate cell stress and damage during sEV production (**Figure 3** and **Supp Table 2**). Analysis of variance underlined that the culture duration significantly affected cell viability (p<0.0001), LDH release (p<0.0001), and CHOP expression (p<0.05) (**Table 2**). The speed of agitation and the cell density significantly increased CHOP expression after longer duration of culture, but had no noticeable effect on cell viability and LDH (**Table 2**). Therefore, in the range of agitation assessed (60 to 120 rpm), there is no optimum to improve cell viability. Increasing the concentration of seeded cells emerged as the only parameter that significantly increased the concentration of sEV produced per mL of culture volume (p<0.0001) (**Figure 3** and **Table 2**).

**Figure 3.**
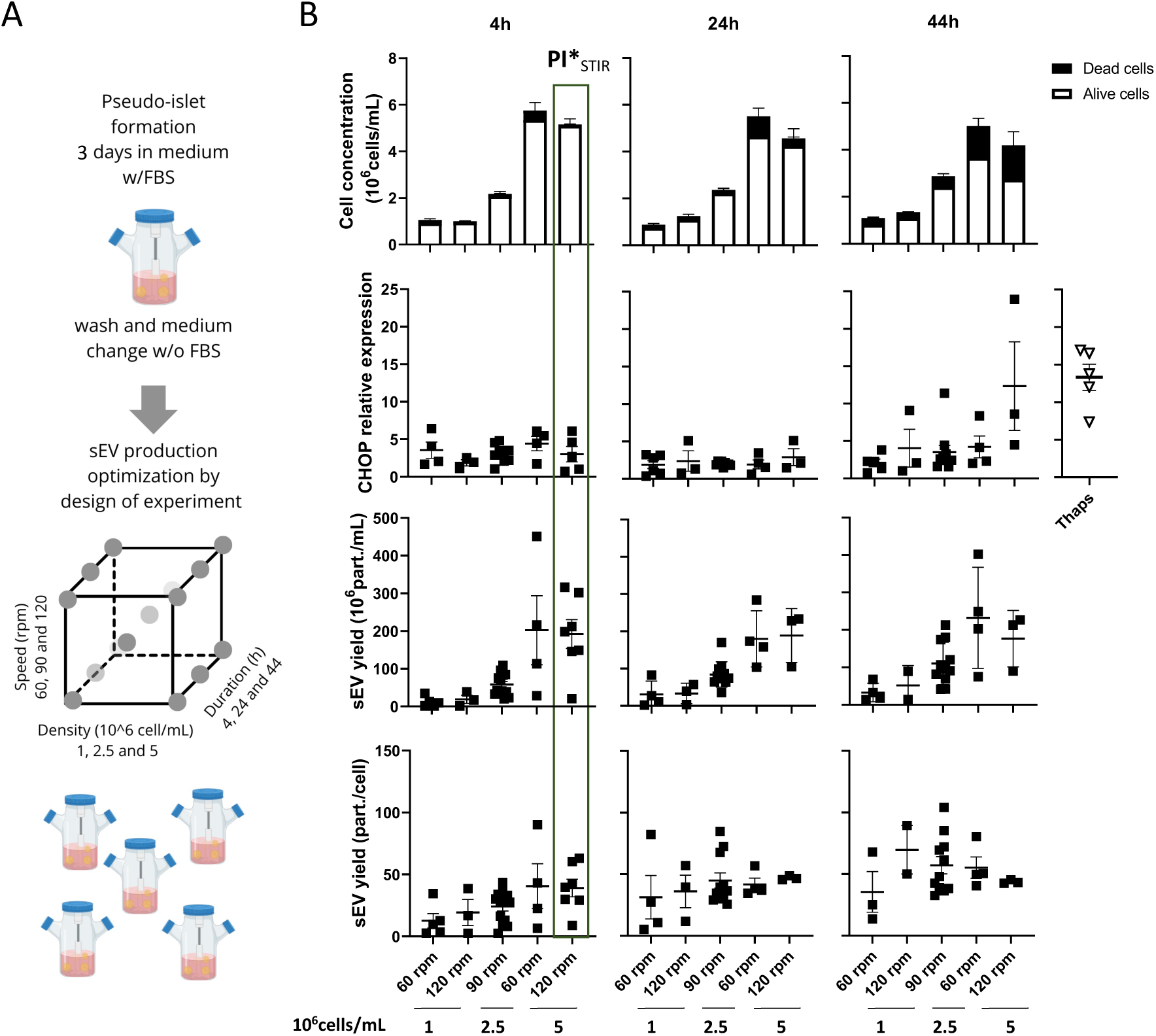
Identification and management of critical cell culturing parameters. (A) Screening design conducted to optimize sEV production by PI cultured in stirred bioreactors. MIN6 PI were seeded at a concentration of 1×10^6^ cell/mL, 2.5×10^6^ cell/mL or 5×10^6^ cell/mL and cultured in a medium without FBS in spinners with stirring at 60, 90 or 120 rpm for 4, 24 or 44 hours. (B) The following response variables were studied: concentration of viable (white) and dead cells (black) (10^6^ cells/mL), relative expression of CHOP compared to PI cultured in spinner in medium w/FBS at 90 rpm, sEV yield per milliliter of culture supernatant or per cell determined by TRPS following sEV precipitation. PI*_STIR_: process settings to improve cell viability, sEV production and mitigate cellular stress responses. Results of screening design are depicted as individual values of biological replicates and means ± SEM from at least three independent experiments.

**Table 2.** Estimated effects of factors, their interactions and the statistical significance thereof to set up process parameters for sEV scalable production.

| Factors | Viability (%) |  | LDH (mUI/10<sup>6</sup> cells) |  | CHOP (relative expression) |  | sEV concentration (10<sup>6</sup> part./mL) |  | sEV yield (part./10<sup>6</sup> cells) |  |
| --- | --- | --- | --- | --- | --- | --- | --- | --- | --- | --- |
|  | effect | p-value | effect | p-value | effect | p-value | effect | p-value | effect | p-value |
| Duration | 19.46 | **<0.0001** | 224.78 | **<0.0001** | 1.84 | **0.0217** | 30.85 | 0.099 | 27.24 | **<0.0001** |
| Speed | 5.67 | 0.082 | -53,24 | 0.3176 | 1.34 | 0.1057 | -3.45 | 0.868 | 7.18 | 0.332 |
| Cell Density | 1.14 | 0.722 | -13.85 | 0.791 | 2.08 | **0.0129** | 166.24 | **<0.0001** | 4.73 | 0.519 |
| Density x Speed | -3.26 | 0.304 | 100.27 | 0.0585 | 1.20 | 0.1461 | -12.51 | 0.545 | -7.86 | 0.282 |
| Speed x Duration | 0.005 | 0.999 | -59.95 | 0.3405 | 3.20 | **0.0119** | -7.36 | 0.768 | 2.66 | 0.765 |
| Density x Duration | -4.48 | 0.231 | 10.13 | 0.8694 | 2.01 | **0.0442** | -13.08 | 0.591 | - 21.05 | **0.016** |
| R<sup>2</sup> | 41.94 % |  | 27,79 % |  | 34,4% |  | 52.8 % |  | 32,2% |  |

\*p-values computed by StatGraphic from the analysis of variance performed for each response reaching statistical significance ≥ 5% are highlighted in **bold**

Indeed, prolonged culture duration allowed increasing the sEV yield per cell (p<0.0001), but globally failed to increase sEV harvest at the end of culture (p=0.0993), due to a strong negative interaction between cell density and culture duration. Based on these results, it appears relevant to produce beta sEV at high cell density and over short periods of culture to preserve cell viability and mitigate cellular stress responses, all while ensuring high sEV production yields. The selected process parameters (PI*) for scalable sEV production were a cell density of 5×10^6^ cell/mL, a culture duration of 4 h and an agitation speed of 120 rpm. Interestingly, these conditions allowed to maintain cell viability at 97.7 ± 1.0 %, with no significant change in the relative expression of CHOP and a 4-fold increase of sEV yield per millilitre of culture volume (p<0.005) in comparison to regular monolayer cultures (**Supp.** Figure 2A-C).

### Impact of the three-step bioprocess on product quality attributes: sEV origin, cargo and immune properties

As EV closely mirror changes in the parental cell, the impact of the optimized production process on sEV morphology (**Figure 4**), origin (**Figure 5**), cargo (**Figure 6**) and immune- modulatory properties in a MLR (**Figure 7**) was analysed. Beta-sEV were isolated and concentrated from culture supernatants by sequential centrifugation, filtration followed by size exclusion chromatography. In some experiments, sEV released by cells exposed to shear stress, as defined by the Kolmogorov scale criteria, were included as a control to study the impact of agitation on sEV quality attributes. To this end, pseudo-islets were agitated at 250 rpm in spinner flasks to achieve a Kolmogorov scale of approximately 36 µm, assuming a power number (Np) of the spinner of 2.51 [55].

**Figure 4.**
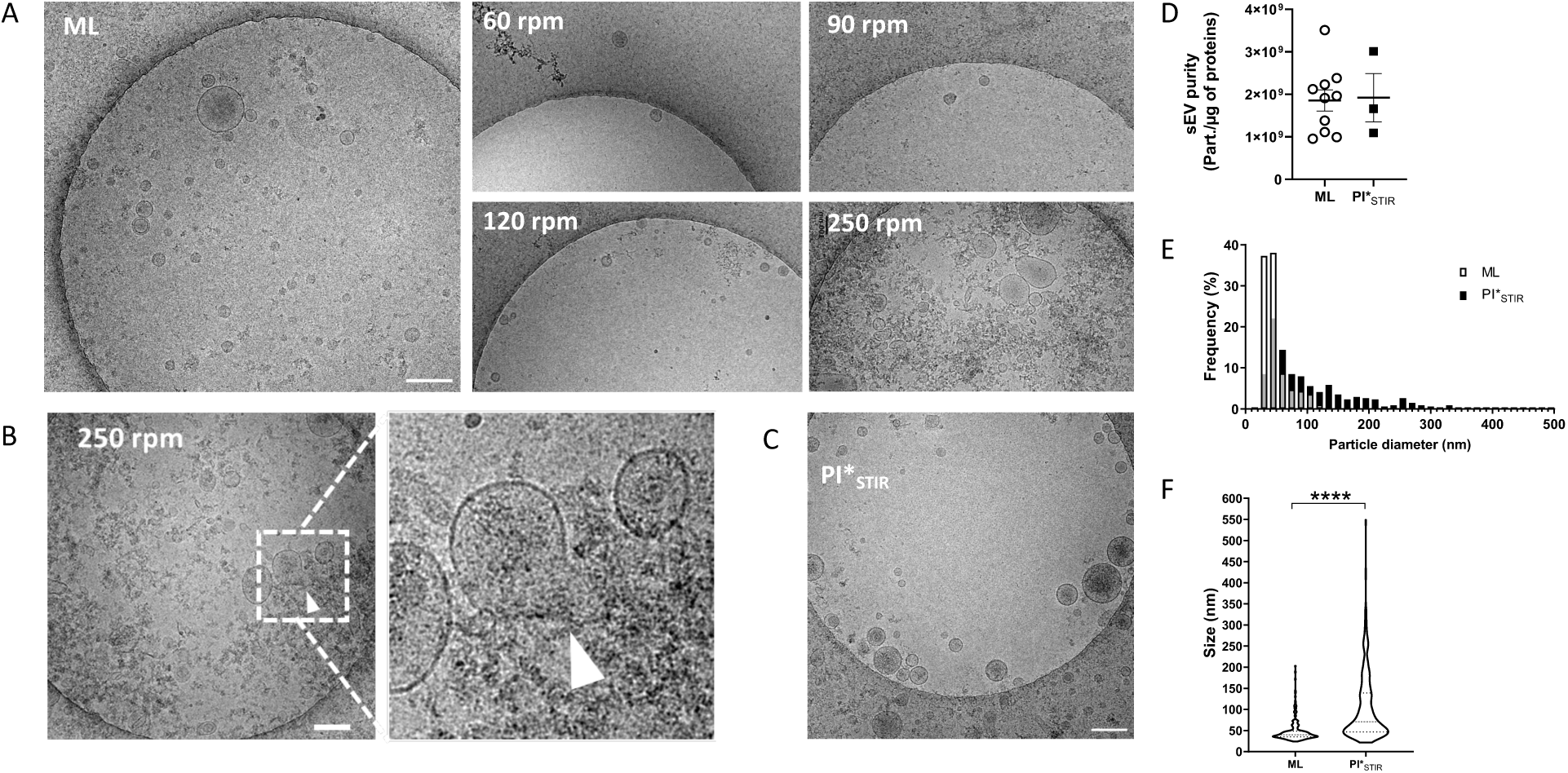
Impact of stirring on sEV morphology and purity. Analyses of beta-sEV purified by TFF-SEC. (A-C) Cryo- electron microscopy images of beta-sEV produced from MIN6 cells (A) grown at medium density (2.5 x 10^6^ cells/mL) as ML or stirred PI at speed as indicated for 24h. (B) under turbulent conditions (250 rpm). Arrows indicate a site of membrane disruption. (C) using PI*_STIR_ settings i.e. density (5 x 10^6^ cells/mL), speed (120 rpm) and culture duration (4h). Scale bar: 200 nm. (D –F) sEV from ML compared to sEV from PI*_STIR_ (D) sEV purity determined by TRPS and Bradford analysis. Results are depicted as individual values and means ± SEM from at least three independent experiments. (E-F) Fiji analysis of sEV size distribution on cryo-electron microscopy images of sEV of ML (n=265) or PI*_STIR_ (n=331). (E) Histogram of relative size frequencies. Bin size: 15 nm. (F) Violin plots of particle size with median and quartiles. Unpaired, parametric t-test (****p<0.0001).

**Figure 5.**
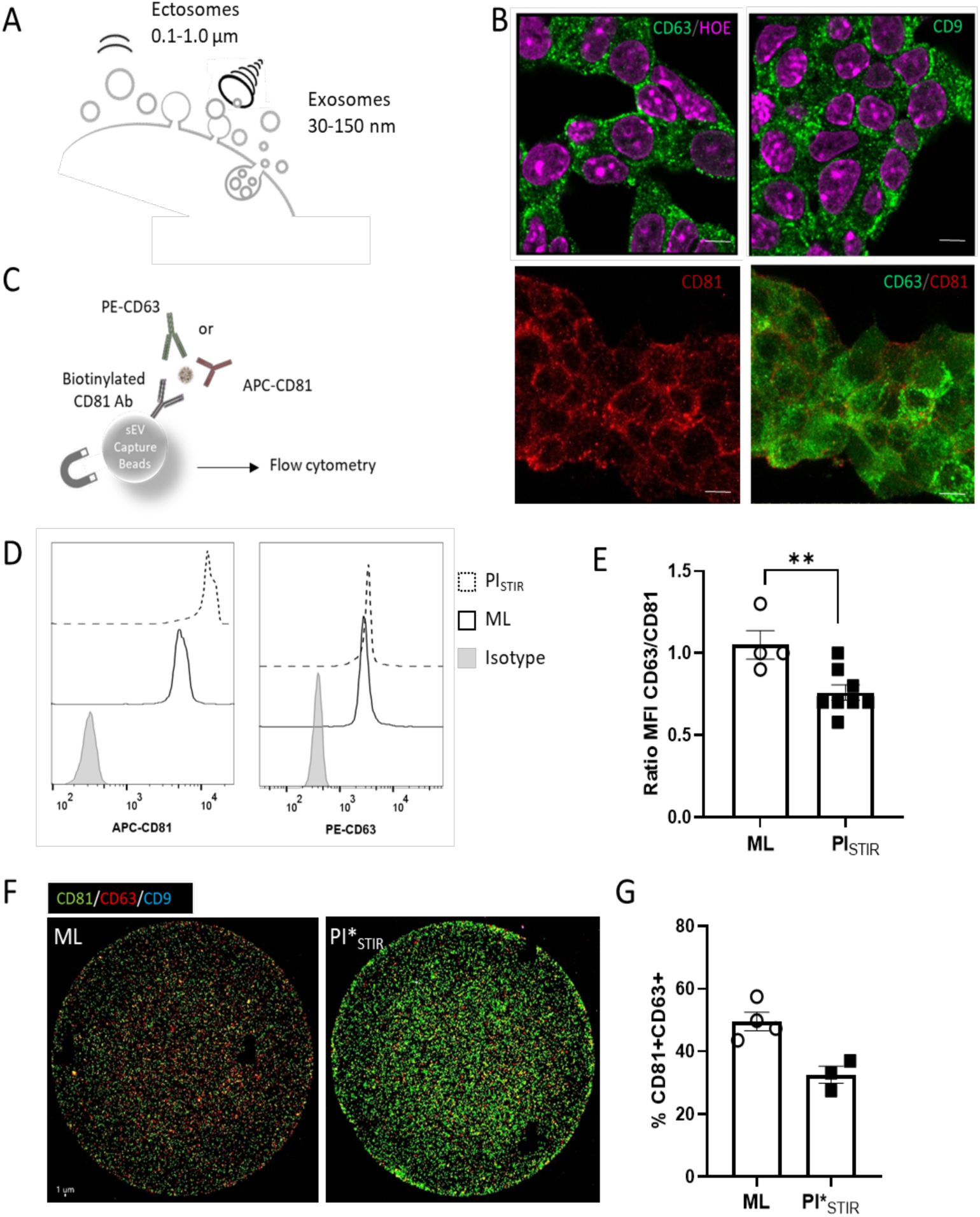
Impact of stirring on sEV biogenesis. Origin of beta-sEV isolated by TFF-SEC from culture supernatants of MIN6 cells grown as static ML or stirred PI_STIR_. (A) Illustration of the hypothetical impact of shear forces on the exosome/ectosome balance. (B) Immunofluorescence staining of CD63 (green), CD9 (green) and CD81 (red) in MIN6 cells with nuclei counterstained with Hoechst (purple). Scale: 5µM (C-D-E) Bead-assisted flow cytometry analysis using anti-CD81 for sEV capture and anti-CD63 (PE) and anti-CD81 (APC) for sEV detection. (D) Representative histograms of CD81 and CD63 mode fluorescence intensities (MFI) on bead-absorbed sEV compared to isotypic controls. (E) MFI CD63/CD81 ratios recorded from four to eight sEV batches derived from ML or PI_STIR_ in three independent experiments Unpaired, parametric t-test (**p<0.01). (F-G) Anti-CD81 immuno- capture on a chip of sEV detected by immunofluorescence with anti-CD81 (green), CD63 (red), CD9 (blue) staining on an Exoview instrument. (G) Percentage of CD63-positive sEV detected following CD81 capture. Results are depicted as individual values and means ± SEM from at least three independent sEV batches.

**Figure 6.**
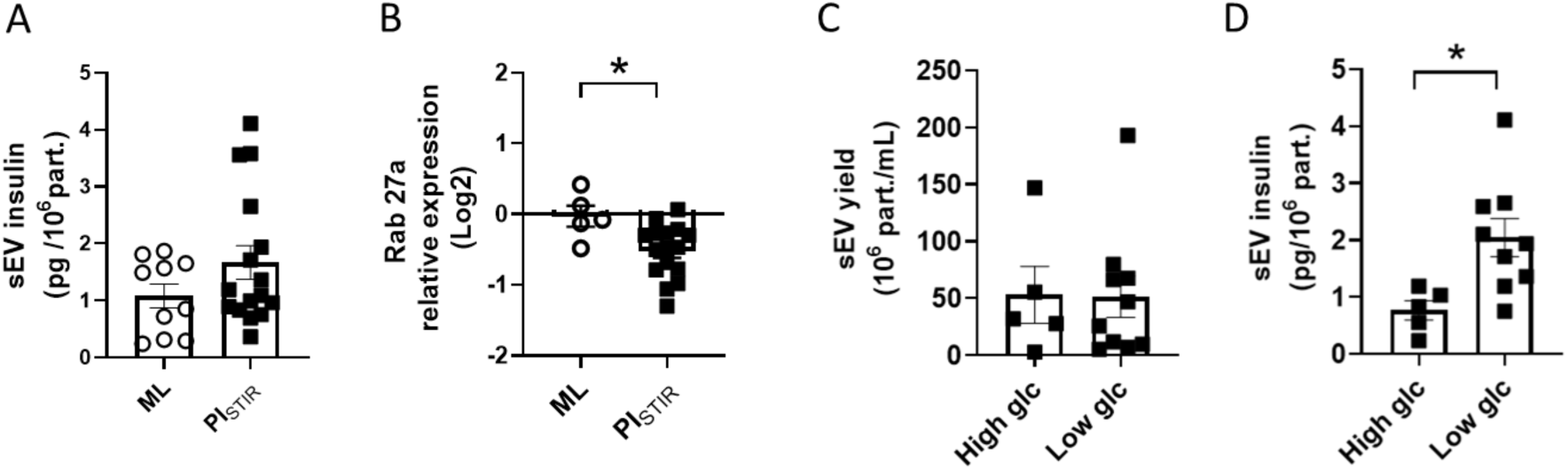
Impact of glucose concentration on beta-sEV insulin cargo. (A-B) MIN6 cells grown as static ML or stirred PI_STIR_. (A) Insulin content determined by ELISA of beta-sEV isolated from culture supernatants after TFF- SEC isolation (B) Relative Rab27a expression in cells represented as fold change of expression with respect to MIN6 ML determined by real-time RT-qPCR. Results from 10 to 15 independent experiments are expressed as mean ± SEM. Unpaired, parametric t-test (*p<0.05). (C-D) Pseudo-islet cultured in DMEM+ 10% FBS in spinners at 90 rpm speed for three days, then, medium was switched to OptiMEM at high [3.5 - 4.5 g/L] or low [1.0 – 1.5 g/L] glucose concentration for 24h prior to sEV isolation by TFF-SEC (C) sEV yield per milliliter of culture supernatant measured by TRPS (D) Insulin quantified by ELISA per 10^6^ particles as determined by TRPS. Results from five to nine independent experiments are expressed as mean ± SEM unpaired, parametric t-test (*p<0.05).

**Figure 7.**
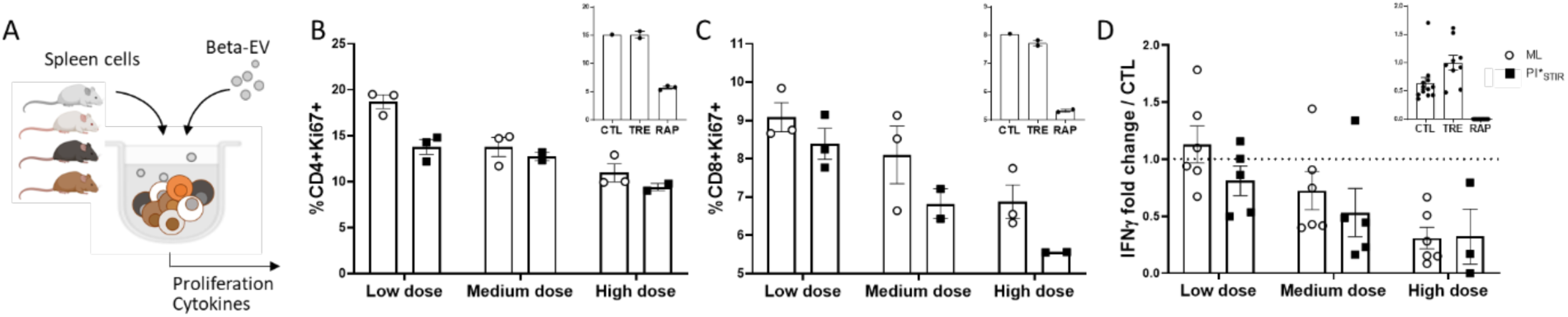
PI*_STIR_ scalable production settings do not alter sEV immune regulatory properties in a MLR *in vitro*. (A) Illustration of a MLR using MHC-mismatched mouse strains (C57BL/6, FVB, BALB/c, C3H). Splenocytes of each strain are mixed and treated with increasing doses of sEV of three different batches derived from ML or PI*_STIR_ cultures: 7 x 10^8^ particles (low dose), 2 x 10^9^ particles (medium dose) and 6 x 10^9^ particles (high dose) in a final volume of 200 µL per well. Rapamycin- (RAPA; 100 nM)) and vehicle only (TRE; PBS 2.5mM trehalose) -treated cells were included as positive and negative controls, respectively. (B and C) After four days of culture, CD4^+^ and CD8^+^ T-lymphocyte proliferation was measured by flow cytometry detection of Ki67. Results from three independent sEV batches are expressed as mean ± SEM. (D) Quantification of IFNlll in culture supernatants expressed as fold change with respect to vehicle only controls. Results from two independent experiments with three independent sEV batches are expressed as mean ± SEM.

In our hands, cryo-EM images of sEV produced under static culture conditions or at low to moderate speed (60 – 120 rpm) presented clear backgrounds (**Figure 4A**). In contrast, sEV produced under strong agitation (250 rpm) presented a dark noisy background due to the presence of amorphous, non-vesicular material. Occasionally, loss of membrane integrity was observed (**Figure 4B**). This was not the case for sEV produced using the improved culture conditions at high-density, 120 rpm for 4 h, as confirmed by cryo-electron microscopy pictures showing numerous vesicles delimited by intact membranes on a clean background (**Figure 4C**). In line with this observation, sEV derived under static or improved stirred conditions were of similar purity as indicated by 1.92 ± 0.57×10^9^ and 1.85 ± 0.25×10^9^ part/µg of protein measured, respectively (**Figure 4D**). Diameter measurement of sEV on high-resolution cryo-electron microscopy images revealed an identical mode size of 45 nm, in the range expected for the size exclusion chromatography column used [35 – 400 nm] (**Figure 4E**). However, sEV produced in static monolayer cultures in comparison to improved conditions presented a shift of the median size from 39.8 nm (24.0 – 179.3) to 70.8 nm (21.5 – 527.9) (median ± range) (**Figure 4F**) providing direct evidence that mechanical agitation influences vesicular morphology (p<0.0001).

Living cells in culture release EV both from the cytoplasmic membrane, so called ectosomes, and vesicles of endocytic origin by fusion of late endosomal compartments containing sEV called exosomes. Ectosomes and exosomes are of overlapping size and similar density and hence not separated by standard isolation techniques. We hypothesized that mechanical forces stretching the cytoplasmic membrane in stirred systems could potentially weigh on the ectosome/exosome balance (**Figure 5A**). Live tracking of intracellular tetraspanin distribution in HeLa cells had revealed earlier preferential mapping of CD63 to endosomal structures in contrast to CD9 and CD81 located essentially at the cytoplasmic membrane [56,57]. In our hands, immunofluorescence staining of MIN6 beta cells confirmed punctuated CD63 staining at intracellular locations, in line with a bona fide exosome marker (**Figure 5B**). Surprisingly, in MIN6 cells, CD9 staining showed similar distribution patterns as CD63, suggesting that cell-type specific differences in tetraspanin distributions might occur. As expected, CD81 spotted mainly to the cytoplasmic membrane. Collectively, this data suggested that CD63 and CD81 staining could be a mean to distinguish between beta sEV of endosomal and ectosomal origin in MIN6 cells. Subsequently, sEV derived from beta cells grown in static monolayer or stirred pseudo-islet cultures were analysed using a bulk bead-assisted flow cytometry assay (**Figure 5C-E**). A ratio of 3 x 10^4^ particles per bead was used to reach signal saturation (data not shown). Our results showed lowered CD63 to CD81 ratio in stirred cultures, suggesting that stirring promotes ectosome rather than exosome release. For sEV derived using the improved culture conditions, immuno-capture-based chip analysis further confirmed lowered percentages of CD63^+^ events at the single EV level: following CD81 immuno-capture, 49% of sEV derived under static conditions proved CD63^+^ against 33% under stirred conditions. The same tendency was observed following CD9 immuno-capture (data not shown) with 66% and 49% of CD63-positive sEV derived under static and stirred conditions, respectively (**Figure 5F- G).**

In pancreatic beta cells, insulin is stored in a pool of secretory granules, out of which 5% are firmly docked to the cytoplasmic membrane and another 20% are located at a distance of less than 200 nm, ready for release by exocytosis upon demand [58]. Under improved stirred conditions, we observed an increase in sEV insulin content from 1.1 ± 0.7 to 1.7 ± 1.2 pg per million of particles (mean ± SEM) in comparison to monolayer cultures, although this difference did not reach statistical significance (**Figure 6**). Evidence had accumulated that members of the small GTPase family Rab27 control fusion of multivesicular bodies to the cytoplasmic membrane and silencing of Rab27 expression reduces exosome release [59–62]. Pancreatic beta cells do not express Rab27b, but Rab27a is present in insulin granules throughout the cytoplasm [63], where it is thought to act as a gatekeeper to prevent spontaneous vesicle fusion in the absence of stimulus-induced Ca^2+^ binding [64,65]. Interestingly, the transcriptomic analysis conducted here revealed a slight though significant 0.7-fold reduction of Rab27 expression in the parental cells under improved stirred conditions (p<0.05) (**Figure 6**), which might constitute an adaptive reaction to counteract insulin efflux due to increased ectosome production. These productions were performed using OptiMEM medium, a low glucose (1 g/L) medium. Further attempts to produce sEV from pseudo-islets cultured in high glucose OptiMEM production medium, revealed a drop of the insulin content in sEV to 0.8 ± 0.2 pg per million of particles (p<0.05), without affecting sEV production yields (**Figure 6**). Altogether, these results suggest that beta cells may evacuate insulin produced in response to high glucose stimulation packed into sEV following nutritional changes in line with the well-described role of EV in waste disposal [66].

Finally, to evaluate the impact of the optimised process on extracellular vesicle immune properties, a MLR was performed. Here, immune cells isolated from spleens of four different mouse strains with major histocompatibility complex-mismatches were mixed *in vitro* triggering robust allogenic immune responses (**Figure 7**). After four days of culture, flow cytometry analysis revealed a dose-dependent reduction of CD4^+^ and CD8^+^ lymphocyte proliferation in the presence of beta-EV derived under static as well as improved stirred conditions. Similarly, lower concentrations of the pro-inflammatory cytokine IFN-γ were measured in culture supernatants illustrating that the aptitude of beta-sEV to dampen T-cell activation was conserved throughout the three-step scalable production workflow.

## Discussion

EV released from healthy mature beta cells have been shown to contribute to beta function and immune homeostasis offering new leverage for diabetes intervention. Given the important quantities of GMP-grade EV required for clinical application, the possibility to produce beta sEV from beta cells grown as pseudo-islets in a scalable STR manufacturing process appears a promising option.

Cultured on hydrophobic surfaces in static or in stirred systems, anchorage-dependent beta cells spontaneously self-aggregate, promoting intercellular interaction. Moderate speed of stirring applied here allows to form small spheroids with a diameter of approximately 60 µm, suited for nutrient and oxygen diffusion into the spheroid core to enhance cell viability [67]. The organization of beta cells into pseudo-islets favours their maturation as evidenced by enhanced expression of the inducible form of the *INS1* gene and the master beta cell transcription factor *NKX6.1* leading to an improved secretory function congruent with previous studies [68–71]. The potential of pseudo-islet culture to promote the differentiation of human embryonic stem cells into pancreatic beta cells has also been explored [72]. More largely, an advantage of 3D culture to produce more physiological sEV from various beta and non-beta cells has been described [70]. Concomitantly, the ability of beta cells to proliferate decreases when grown as pseudo-islets, in line with the fundamental inverse relationship between cell proliferation and differentiation, thoroughly reviewed elsewhere [73]. In the same way, immature primary beta cells isolated from newborn pigs are able to proliferate, a capacity lost upon maturation in islets from adult pigs [74]. Therefore, in order to produce large amounts of sEV from mature cells, a three-phase upstream process is required, with culture conditions favouring first cell proliferation in monolayers (phase I), followed by differentiation in pseudo-islets (phase II), prior to production (phase III).

A major challenge in EV production is the requirement to work with EV-free, preferentially chemically defined media, that may be source of cellular alterations and, consequently, affect sEV quality attributes. In our settings, serum starvation over a short period (24 h) does not reduce cell viability in static monolayer and spheroid suspension culture. However, stirring triggers cellular stress responses and decreases viability. As a general rule, for a turbulent vortex to cause local dissipation of energy triggering cell death, the Kolmogorov scale must be smaller than or similar to the size of the cells. In this study, after three days of formation, pseudo-islets had a mean diameter of approx. 60 µm and were stirred at 60, 90 or 120 rpm, representing a Kolmogorov scale of up to 100 µm [75]. On this basis, stirring up to 120 rpm should allow to avoid shear stress damage during the production phase. Using a DoE approach, the process parameters are adjusted by reducing culture duration and increasing cell density at seeding in order to promote cell viability and sEV production yields. These settings manage to sustain a cell viability higher than 90% and to avoid up-regulation of markers of stress in MIN6 spheroid culture. While sEV concentration markedly increases with cell density at short culture duration, surprisingly, no significant accumulation of the sEV produced occurred over time suggesting their reabsorption by the cells leading to an equilibrium between uptake and release. Therefore, to increase production yield, our results emphasize the value of developing EV production processes operating in perfused or simple continuous mode, which would allow for the regular collection of sEV [76]. In addition, replenishment of nutrients and removal of by-products in the perfusion mode should further be a means to reach even higher cell densities and sEV yields. To do so, the main challenge would be to formulate chemically defined media able to sustain cell viability, growth and maturation for long periods, compatible with downstream processes.

In comparison to the monolayer, no decrease in the yield of sEV produced per cell is observed in 3D culture, despite an estimated 7-fold reduction of the cell surface exchange area. The cell specific productivity even increases when sEV are produced under stirred conditions. Indeed, it is a well-established fact that physical stimulation as exerted by shearing forces enhances vesiculation by applying tension to the cytoplasmic membrane [77]. Here, cryo-electron microscopy analysis confirms the absence of amorphous, non-vesicular material at low to moderate speed of agitation in contrast to high-speed controls. More importantly, we report for the first time that a higher stirring speed induce loss of sEV membrane integrity and contamination of the final product by non-vesicular material comprising an inherent potential risk of sEV cargo and function loss. Process set up at moderate speed allows to warrant high purity and integrity of sEV produced. Yet, despite size selection throughout the downstream purification process, sEV derived under optimised condition present a subtle increase of the median size suggesting mechanical reinforcement of membrane budding, exerted on the cytoplasmic membrane. The mode size remained unchanged.

Downstream purification processes commonly lead to the isolation of sEV of heterogeneous subcellular origin and function. This study uses EV surface marker analysis as a new means to investigate on the implications of the upstream processes on the genesis of sEV produced. In contrast to earlier, sometimes discrepant observations [56,78,79], CD9 does not appear a reliable ectosome marker in MIN6 cells in our hands. Indeed, beta cells are highly specialized secretory cells, a function intimately linked to endo-exocytotic membrane trafficking, which might affect CD9 distribution. Furthermore, single-cell RNA-seq has revealed that CD9 is a negative marker of glucose-responsive pancreatic beta-like cells derived from human pluripotent stem cells, in line with a distinct role for CD9 in beta cells [80]. As expected, CD81 and CD63 staining in MIN6 cells mapped preferentially to cytoplasmic membrane and intracellular compartments and were used as ecto-and exosome markers, respectively. The lowered CD63/CD81 ratios observed in stirred systems here present a first line of evidence that more ectosomes are produced, including at speed below standard turbulence thresholds.

In view of this finding, we hypothesized that EV production in stirred systems may promote the incorporation of cytosolic rather than endosomal components into cytoplasmic membrane vehicles with distinct cargo. In beta cells, massive amounts of ready-to-release insulin are stored inside secretory granules close to or docked to the cytoplasmic membrane [58]. ELISA quantification shows an upward trend of the insulin export in sEV derived under improved stirred conditions in comparison to sEV from static monolayer culture. This trend does not reach significance, which might be due to the average size 300 nm of mature secretory granules, not retained in the sEV fraction recovered by the downstream process [81]. Notwithstanding the artificial nature and its potential hidden drawbacks, formation of sEV under stirring may be of particular benefit for ectosome-enriched compound-loading e.g. cytosolic or membrane-bound substances for therapy or vaccination.

We further queried whether glucose stimulation during the production phase could enhance export of insulin inside beta-sEV. Surprisingly, we observed a reduction of insulin content in our beta-sEV when produced in high glucose (25 mM) production medium, compared to low glucose (5.5 mM) medium. A plausible explanation might be that beta cells export excessive insulin inside EV in adaption to changes from high to low glucose environments [66]. More largely, multi-step bioprocesses combining a phase of culture triggering cargo biogenesis and a phase of EV production with stimulant withdrawal e.g. starvation media could proof useful to direct EV composition.

Maintenance of plasma membrane integrity is essential for cell survival and function. It is tempting to speculate that mechanistic stress and loss of cellular material trigger resistance and membrane repair mechanisms. Among key regulators of EV release, Rab27a, a member of the Ras-associated protein GTPase family, drives the fusion of late endosomal compartments with the cytoplasmic membrane, leading to exosome secretion [82]. Knock- down studies further confirm Rab27a-dependence of exosome release *in vitro* and *in vivo* (reviewed in [83]). Ectopic expression of EGFP-Rab27a fusion proteins in transgenic mice reveals highest expression in specialized secretory cells performing regulated exocytosis including both alpha and beta cells of the endocrine pancreas [84]. A plausible explanation for the drop in Rab27a expression apparent at mild stirring in our optimised process might be the activation of compensatory mechanism to reduce material disposal via secretory pathways.

Assuming that the biological activity of sEV batches results from the sum of the activity of EV subpopulations composing the batch [85], we next sought to instruct on the impact of phenotypic changes induced by stirring on the beta sEV immune-modulatory capacity. Pre-existing potency assays developed for drug screening in transplantation research have been tested essentially for sEV derived from mesenchymal stem cell (MSC). Yet, these tests still require standardisation and proper reference material and the overall predictive potential of therapeutic efficacy remains low [86]. Comparative studies on MSC sEV highlight the correlation of *in vitro* allogenic MLR to *in vivo* efficiency in graft versus host disease models [87]. In an in-house developed murine MLR, sEV produced using the optimised process maintained their ability to reduce T-lymphocyte proliferation and IFN-γ cytokine secretion in a dose-dependent manner. Though encouraging, this overall anti-inflammatory activity calls for confirmation in a T1D antigen-specific reaction. In pursuit of this work, we aim to identify the quality attributes responsible for beta sEV therapeutic activity in mouse models of the disease required to streamline relevant potency assays.

In conclusion, this study demonstrates the feasibility to produce sEV from mature stress-sensitive beta cells cultured as small pseudo-islets in STR, amenable to upscaling. In perspective, an interesting solution to obtain a full-scale process, could be the use of microporous microcarriers for beta cell expansion, which offer the possibility to culture anchorage-dependent cells as monolayers on spherical surfaces that can be grown in suspension [88]. Interestingly, the production of sEV in STR under starving conditions may be of particular benefit for ectosome-enriched compound-loading e.g. cytosolic or membrane-bound substances for therapy or vaccination.

## Acknowledgments

We thank the French National Research Agency (ANR-21-CE18-0036, Bioprod-sEVforT1D) for funding the project and the Ministry of Agriculture and Food Sovereignty and the Pays de la Loire region (CPER #00153032) for supporting the PhD of Thibaud Dauphin. We are also grateful to Eugénie Lahet, Claire Boursier, Mathilde Laubert, Célia Bonnouvrier, Mailys Le Devehat, Lucie Grare and Quentin Le Yondre for their contribution to data collection as part of their training or internships.

## Conflict of interest

None of the authors reported a potential conflict of interest.

## Author contributions

L.dB., J-M.B., S.B. and M.M. conceived the study. L.dB., T.D., Q.P., A.D., L.D., D.J., S.B. and M.M. performed the experiments. L.dB., T.D., Q.P., L.D., G.M., B.M., A.S., S.B. and M.M. analysed the data. L.dB., S.B. and M.M. wrote the manuscript. J-M.B., B.L. and J.H. reviewed and edited the manuscript.

## Notes

### Competing Interest Statement

The authors have declared no competing interest.

