## Supplemental for "Three-step scalable production of extracellular vesicles from pancreatic beta cells in stirred tank bioreactors promotes cell maturation and release of ectosomes with preserved immunomodulatory properties"

### Supplemental figures

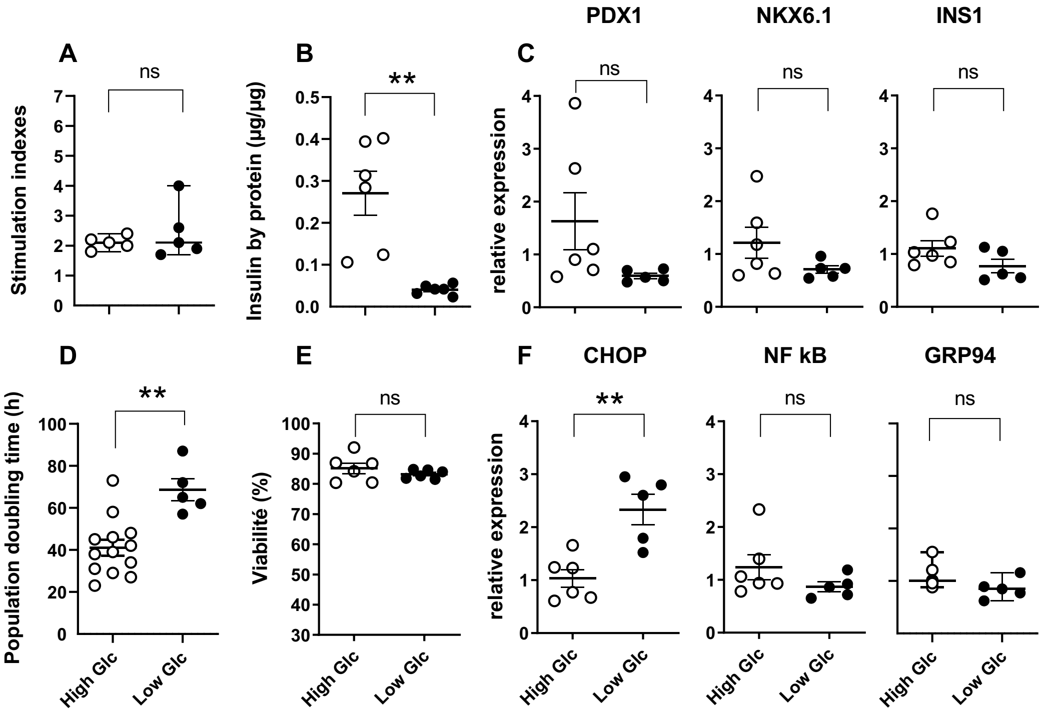


**Supp. Figure 1.** Influence of glucose concentration on MIN6 cell phenotype during the amplification step in monolayer culture. MIN6 cells were cultured in DMEM + 10% FBS at high [3.5 - 4.5 g/L] or low [1.0 – 1.5 g/L] glucose concentration. Glucose concentration was daily-assessed (BioProfile, Nova Biomedical) and adjusted whenever required to reach the targeted ranges. The cells were grown for one week to determine the population doubling time. Beta cell viability and function analysis was performed on day 3. (A) Stimulation index (ratio of insulin after high glucose plus theophylline stimulation over basal insulin secretion release, mM/mM) (B) Intracellular insulin per total protein content (µg insulin/µg protein). (C) Relative quantitative RT-PCR analysis of expression of beta cell transcription factors PDX1, NK6 homeobox 1 (NKX6.1) and INS1. (D) Population doubling time (hour) calculated during the exponential growth phase. (E) Cell viability calculated as the percentage of trypan blue negative viable cells out of total cells. Dead and lysed cells in cell supernatants were estimated by the LDH assay. (F) Relative quantitative RT-PCR expression analysis of CHOP, nuclear factor kappa-light-enhancer of activated B cells *(NF*-kB*)* and glucose reactive protein 94 *(*GRP94*)*. Results from at least five independent experiments are expressed as mean ± SEM, unpaired, parametric t-test (**p<0.01).

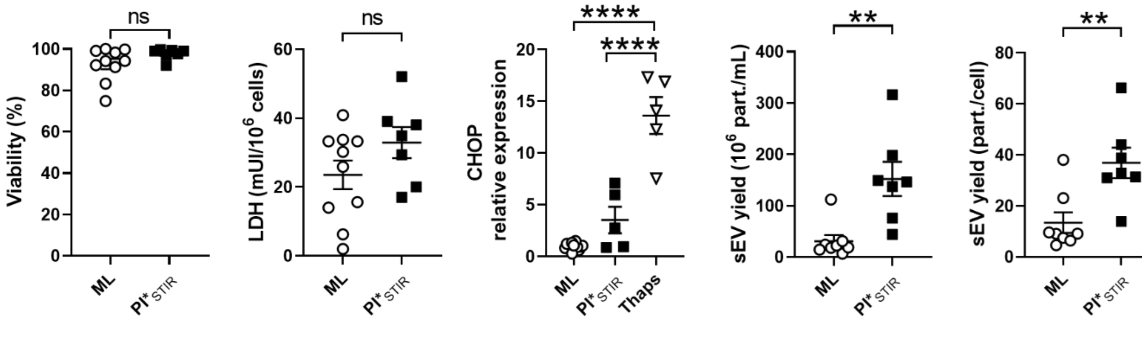


**Supp. Figure** ***2.*** Validation of optimized process (PI*$_{STIR}$) compared to monolayer culture process (ML) (A) Cell viability was assessed by measuring the total of viable cells after trypsin and blue trypan staining and of dead cells by measuring the LDH activity in supernatant. (B) Relative CHOP expression of MIN6 cell compared to ML and ML treated with 1µM of thapsigargin (Thaps) during 24h in medium w/FBS. (C) Concentration and yield of sEV isolated by precipitation method. Results from at least five independent experiments are expressed as mean ± SEM*,* unpaired, parametric t-test (*$**p<0.01$, $****p<0.0001$).*

### Supplemental table

**Supp. Table 1.** Factorial design matrix and experimental results obtained for the screening DoE

| Run | Factors | | | Responses | | | |
|---|---|---|---|---|---|---|---|
| | Culture mode | Hydrodynamic | Medium | Viability (%) | LDH (mUI/$10^6$ cells ) | CHOP (relative expression) | sEV ($10^6$part./mL) |
| **1** | ML | static | w/ FBS | 94.6 ± 3.4 | 8.6 ± 5.5 | 0.6± 0.1 | [redacted] |
| **2** | PI | static | w/ FBS | 97.4 ± 1.0 | 23.2 ± 7.0 | 1.90 ± 0.5 | [redacted] |
| **3** | PI | stirred | w/ FBS | 88.7 ± 10.6 | 29.6 ± 16.0 | 6.5 ± 2.1 | [redacted] |
| **4** | ML | static | w/o FBS | 93.3 ± 5.0 | 22.0 ± 11.1 | 1.5 ± 0.4 | 10.2 ± 7.34 |
| **5** | PI | static | w/o FBS | 93.8 ± 4.2 | 88.1 ± 29.9 | 2.4 ± 0.8 | 1.7 ± 1.0 |
| **6** | PI | stirred | w/o FBS | 92.1 ± 5.4 | 57.3 ± 30.2 | 5.4 ± 3.1 | 38.5 ± 17.5 |

**Mean and SEM of at least five independent experiments*

**Supp. Table 2.** Factorial design matrix and experimental results obtained for the response to surface DoE.

| Run | Factors | | | Responses* | | | |
|---|---|---|---|---|---|---|---|
| | Speed (rpm) | Density ($10^6$ cells/mL) | Duration (h) | Viability (%) | LDH (mUI/$10^6$ cells) | CHOP (relative expression) | sEV concentration ($10^6$ part./mL) |
| **1** | 60 | 1 | 4 | 91.6 ± 6.9 | 32.5 ± 15.1 | 3.5 ± 1.7 | 12.0 ± 9.3 |
| **2** | 120 | 1 | 4 | 94.9 ± 2.6 | 29.2 ± 13.4 | 1.8 ± 0.5 | 18.9 ± 13.1 |
| **3** | 60 | 5 | 4 | 93.7 ± 6.9 | 29.5 ± 10.8 | 4.4 ± 1.4 | 202.2 ± 131.4 |
| **4** | 120 | 5 | 4 | 97.9 ± 1.5 | 35.3 ± 10.2 | 3.0 ± 1.9 | 191.9 ± 74.3 |
| **5** | 90 | 2.5 | 4 | 96.5 ± 2.3 | 37.1 ± 16.1 | 3.1 ± 0.9 | 57.9 ± 26.3 |
| **1** | 60 | 1 | 24 | 85.1 ± 43.8 | 119.2 ± 77.5 | 1.9 ± 1.0 | 30.6 ± 25.8 |
| **2** | 120 | 1 | 24 | 88.5 ± 6.8 | 55.7 ± 34.1 | 2.4 ± 1.8 | 33.3 ± 19.9 |
| **3** | 60 | 5 | 24 | 84.9 ± 7.5 | 66.63 ± 14.1 | 1.9 ± 0.9 | 179.8 ± 52.1 |
| **4** | 120 | 5 | 24 | 92.6 ± 2.2 | 108 ± 21.2 | 2.9 ± 1.5 | 188.7± 56.0 |
| **5** | 90 | 2.5 | 24 | 94.3 ± 5.6 | 71.1± 38.0 | 2.01 ± 0.2 | 84.1 ± 22.3 |
| **1** | 60 | 1 | 44 | 64.4 ± 18.4 | 278.1 ± 566.5 | 2.1 ± 0.8 | 32.9 ± 17.5 |
| **2** | 120 | 1 | 44 | 80.9 ± 3.4 | 126.5 ± 106.8 | 4.1 ± 3.3 | 51.9 ± 37.6 |
| **3** | 60 | 5 | 44 | 72.4 ± 13.0 | 231.8 ± 68.9 | 4.2 ± 2.1 | 232.3 ± 93.2 |
| **4** | 120 | 5 | 44 | 64.8 ± 20.0 | 449.5 ± 123.1 | 12.3± 7.7 | 176.7 ± 57.8 |
| **5** | 90 | 2.5 | 44 | 85.1 ± 11.2 | 131.4 ± 80.1 | 3.25 ± 1.8 | 110.6 ± 44.5 |

*Mean and SEM of at least three independent experiments*
